# Long-term neuron tracking reveals balance of stability and plasticity in functional properties

**DOI:** 10.1101/2025.03.14.643204

**Authors:** Hung-Yun Lu, Hannah M. Stealey, Yi Zhao, Cole R. Barnett, Enrique Contreras-Hernandez, Samantha R. Santacruz

## Abstract

Neural stability is essential for executing learned motor behaviors while plasticity provides the flexibility needed to adapt to new tasks and environments. Although low-dimensional neural population dynamics exhibit long-term stability, the extent to which individual neurons retain their functional properties over time and balance the need for both stability and plasticity remains an open question. Tracking individual neurons across multiple recording sessions is crucial to addressing this question, yet conventional methods face challenges such as electrode drift, waveform variability, and large inter-electrode distances that limit the number of channels a neuron is observed on. Here, we introduce a waveform-based neuron tracking method optimized for standard microelectrode arrays, enabling the identification of the same neurons across sessions without relying on spatial overlap, a strategy commonly leveraged with high-density electrode arrays. We apply this method to assess the longitudinal stability of multiple neural properties, including firing rates, inter-spike intervals, tuning properties, and spike-field interactions. Our findings reveal that while spike waveform properties remain stable, certain functional properties such as ISI and tuning can exhibit gradual shifts, suggesting a balance between neural stability and plasticity. Understanding the persistence of individual neural signals provides insight into learning and adaptation while advancing the study of neural stability and plasticity over extended timescales. Beyond basic neuroscience, this framework has potential to enhance the long-term reliability of brain-machine interfaces and closed-loop deep brain stimulation systems that rely on chronic neural sensing.

## Introduction

Neural circuits exhibit remarkable stability in generating consistent activity patterns while retaining the flexibility to adapt to new conditions, learning processes, and environmental changes (1–4). Despite biological variability at multiple levels of neural organization, from ion channels (5), synapses (6), to neurons (7), the brain maintains reliable representations that support stable cognition, perception, and behavior. However, it remains unclear whether this stability arises from stable underlying neural dynamics or whether individual neural signals fluctuate over time. Previous research has demonstrated that latent neural dynamics, or low-dimensional population representations, exhibit long-term stability (8–10), yet the degree to which individual neurons maintain stable firing patterns across sessions remains unresolved.

The question of neural stability has been the subject of ongoing debate. Some studies report that individual neurons retain highly stable functional properties (3,11–13), whereas others suggest substantial variability in neural activity over time (14–17). A recent study examined long-term neural recordings and found that individual neurons appeared stable, attributing fluctuations primarily to unrestricted kinematics (18). However, their study did not fully characterize other aspects of neural stability, such as spiking dynamics, tuning properties, and interactions with local field potentials. Additionally, their dataset covered a relatively short time window, leaving open the question of whether individual neural properties remain stable over longer timescales.

A major challenge in addressing this question is the difficulty of reliably identifying the same neurons across multiple electrophysiological recording sessions (2,3,19,20). Chronic electrode arrays enable long-term neural recordings and have been widely used to study behavior and learning over months and years (19–21). While these arrays allow for recordings from consistent brain regions, factors such as brain micromovements and probe displacement (22,23) introduce uncertainty in determining whether a neuron recorded on the same electrode channel across sessions is truly the same neuron. Despite these limitations, chronic microelectrode arrays remain the best available method across species for tracking neural activity longitudinally. However, a reliable computational framework is necessary to track neurons effectively across sessions of data from long-term studies.

Several approaches have been developed to address this problem (3,11,14,18,24–30). Early methods relied on qualitative waveform comparisons to match neurons across sessions (1,3,11). More recently, algorithms, such as UnitMatch (29) and Earth Mover’s Distance (30), have been introduced to improve longitudinal neuron tracking. However, many methods were designed for high-density probes (29–31) or tetrode arrays (25,28). In these cases, they specifically dealt with spatial overlaps, where neurons could often be detected simultaneously by multiple channels due to the short inter-electrode distances (for example, 25 µm center-to-center spacing in Neuropixels, (32)). In contrast, conventional microelectrode arrays, such as Utah arrays (33,34), typically feature large inter-electrode distances (> 200 µm, (27)) that result in a lack of such spatial redundancy, making waveform-based tracking a more viable approach (26). Despite the widespread use of conventional microelectrode arrays in neurophysiological research, robust tracking methods tailored to their unique recording characteristics remain underexplored.

In this study, we introduce a waveform-based neuron tracking method optimized for conventional microelectrode arrays, enabling the identification of the same neurons across multiple sessions from long-term (3+ months) experiments without accounting for spatial oversampling across channels. Using this approach, we systematically assess the stability of individual neurons by analyzing their firing rates, inter-spike intervals (ISI), tuning properties, and spike-field relationships (35,36). Our analysis focuses on spiking activity recorded from motor areas, providing a comprehensive evaluation of how individual neurons contribute to persistent motor representations over time. However, this framework is broadly applicable to spiking activity recorded longitudinally from any brain area, making it a valuable tool for investigating neural stability across different circuits.

The ability to track neurons across sessions has significant implications for both basic neuroscience and applied neurotechnology, such brain-machine interface (BMI) research (37) and deep brain stimulation (38). From a theoretical perspective, identifying stable neurons enhances our understanding of long-term neural representations and the plasticity mechanisms underlying motor learning (10). From an applied perspective, neuron tracking improves the reliability of chronic neural interfaces. For example, for BMIs this can enable the selection of stable units for decoding, thereby enhancing long-term performance (3). By developing and validating a framework for multi-session neuron tracking, this study provides a foundation for future research into the persistence and adaptability of neural circuits in the motor cortex.

## Materials and Methods

### Experiments

Two male rhesus macaques (Macaca mulatta) participated in this study (Monkey A: age 5 years, ∼9.5 kg; Monkey B: age 5 years, ∼8.8 kg). All experimental procedures were approved by the Institutional Animal Care and Use Committee (IACUC) of The University of Texas at Austin. The animals were housed in a controlled environment with a 12-hour light-dark cycle, regulated temperature, and humidity. They were socially housed whenever possible to promote natural interactions, with access to species-appropriate environmental enrichment, including foraging opportunities, climbing structures, and novel objects to encourage cognitive engagement. The primates were fed a balanced diet with water available ad libitum. Prior to training, fluids will gradually be reduced in order to regulate fluid intake and ultimately use fluids as a motivating factor in trained behaviors. Body weight and overall health were closely monitored by veterinary staff, and all animals received regular health assessments.

Each monkey was unilaterally implanted with tungsten microelectrode arrays (MEAs) targeting the primary motor cortex (M1) and premotor cortex (PMd). Monkey A was implanted with a 64-channel MEA, while Monkey B received a 128-channel MEA. The implantation sites were determined using stereotaxic coordinates, pre-operative magnetic resonance images, and cortical landmarks.

Experiments and training are typically performed five days per week, for an average of 3-6 hours/day and will always comply with the published guidelines to promote the psychological well-being of non-human primates. The subjects were trained to perform a center-out motor task under brain-machine interface (BMI) control using a biomimetic decoder (Fig 1A). Behavioral metrics and neural signals, including spiking activity and local field potentials (LFP), were recorded simultaneously. Neural data was acquired using the Grapevine NIP electrophysiology system (Ripple Neuro, Salt Lake City, UT). Before each session, spike sorting was performed online using the hoop method (39) in the Grapevine NIP system. In this study, we defined each source of neural activities as a “sorted neuron” (see Glossary below). A subset of sorted neurons was used to control the cursor via a Kalman filter (40–42).

**Fig 1.**
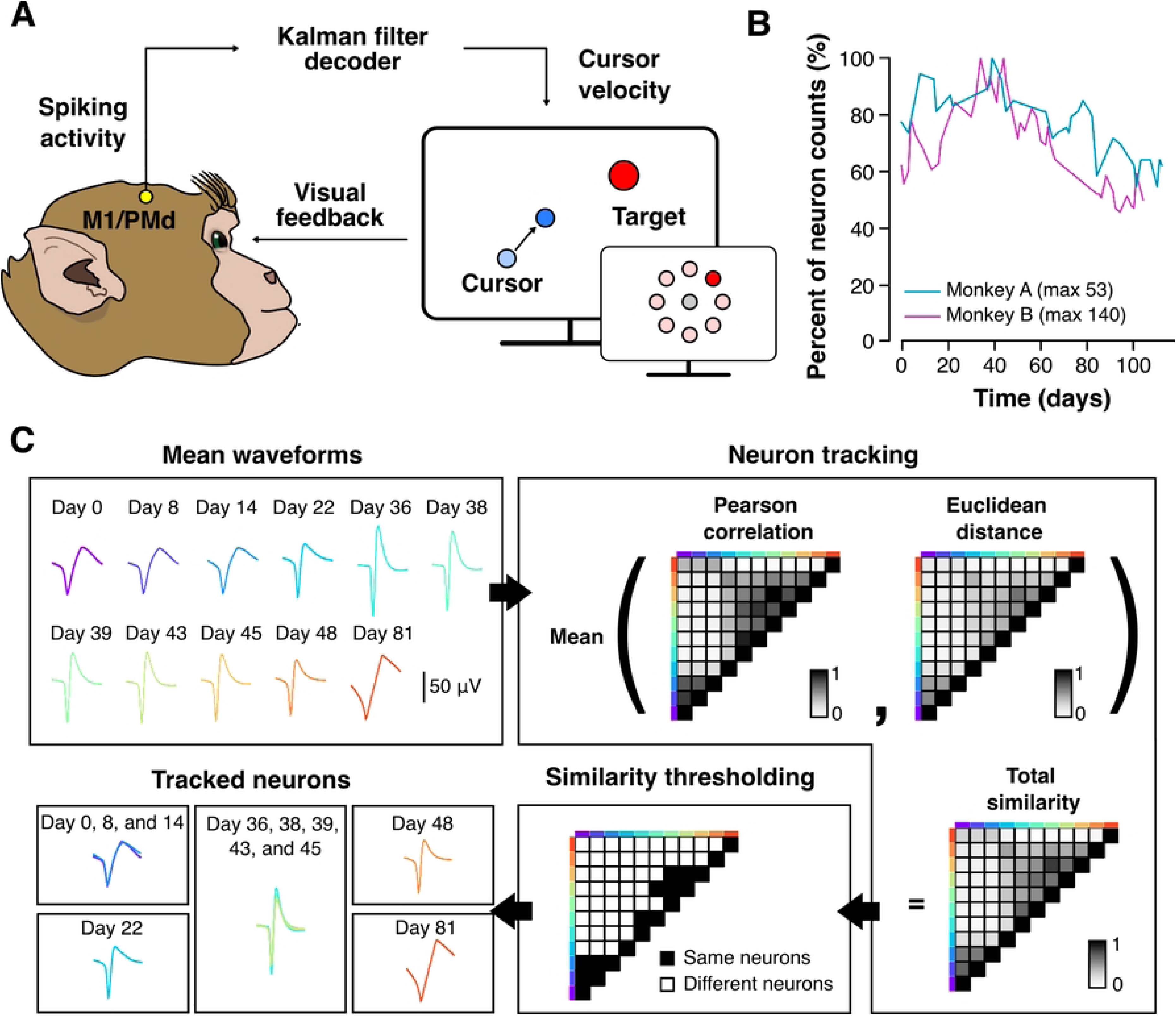
Experimental setup and neuron tracking algorithm. (A) The center-out task required the subjects to control the cursor through brain-machine interface in order to hit one of the eight peripheral targets from the center. The spiking activities of the motor regions (M1 and PMd) were decoded by a Kalman filter into instantaneous cursor velocities. (B) The percentage of neuron counts over time. Each point represents the total number of sorted neurons divided by the maximal number of sorted neurons of the subject (Monkey A: 53 neurons; Monkey B: 140 neurons). (C) The eleven mean waveforms (MWs) of an example channel from Monkey A were used to demonstrate the neuron tracking pipeline. The total similarity matrix (shape of 11 by 11; only upper triangular matrices was shown) was the average of two rescaled metrics, including Fisher-transformed Pearson correlation and log-transformed Euclidean distance. The total similarity matrix was examined by a similarity threshold that was estimated from MWs that were known different. A sorted neuron was included in a tracked neuron if it had a similarity score above the threshold with at least one sorted neuron in the group. By the end of this pipeline, we obtained five tracked neurons. The tracked neurons that had less than three sessions (Day 22, Day 48, and Day 81) were not used for further analyses. The other tracked neurons (Day 0, 8, and 14 and Day 36, 38, 39, 43, and 45) had a duration of 15 and 10 days, respectively.

In each trial, the monkey was required to hold the cursor at the center of the screen for 200 msec before a peripheral target appeared in one of eight possible locations radially distributed around the center of the screen. The subject then moved the cursor toward the peripheral target and held it there for 200 msec within a 10-second window to successfully complete the trial. Each session included in the analysis consisted of 336 successful trials, with 42 trials per target direction. Only neurons with an average firing rate greater than 1 Hz and peak-to-trough amplitude greater than 80 µV were included in the analysis to ensure sufficient data quality. In total, Monkey A completed 34 sessions over 113 days, while Monkey B completed 44 sessions over 105 days (Fig 1B).

### Neuron tracking algorithm

To track neurons across sessions, we assumed that neurons could not migrate between electrodes due to the large inter-electrode distances (26,27) in our microelectrode arrays (500-750 µm). Therefore, tracking was performed separately within each channel based on the mean waveforms (MW) extracted from individual online sorted units from each session.

The tracking algorithm consisted of three main steps: (1) computing pairwise MW similarities between spiking activity from pairs of sessions, (2) defining a threshold similarity score, and (3) clustering neurons into longitudinally tracked groups (Fig 1C). Channels with fewer than three sorted neurons across all sessions were excluded to ensure sufficient data for meaningful tracking. Since multiple neurons could be sorted from a single channel within a session, our algorithm treated all sorted neurons within the same channel as candidates for longitudinal tracking, without enforcing constraints on whether multiple sorted neurons in the same session should be clustered into a single tracked neuron. If two or more sorted neurons from the same session were mistakenly grouped together, we resolved these conflicts post-hoc. This approach ensured that tracking decisions were not affected by inconsistencies in indexing of sorted neurons on the same channel across sessions, such as when the same neuron was assigned a different index on different sessions.

### Pairwise MW similarities

For most electrode channels with spiking activity, only one online sorted neuron was identified, resulting in a single MW from the neuron. For channels with more than one online sorted neurons, a separate MW was derived for each corresponding online sorted neuron. Neuron tracking was performed per channel based solely on MWs from online sorted neurons from each session. Specifically, all MWs from each channel were compared across sessions. Pearson correlation and Euclidean distance were used to quantify similarity, as they capture different aspects of waveform properties (24,29). Pearson correlations were Fisher-transformed, and Euclidean distances were log-transformed to normalize each distribution. These transformed similarity scores were then rescaled, where one represented the highest similarity and zero represented the lowest. The final total similarity score was computed as the average of the rescaled correlation and distance metrics. The resulting similarity matrices were stored alongside their corresponding channel and unit information for downstream analysis.

### Defining a threshold similarity score

To determine the similarity threshold for neuron tracking, we generated null distributions of similarity scores from MW pairs known to belong to different neurons (25). Specifically, MWs were randomly paired from distinct channels across sessions in batches, ensuring that the number of MWs in each batch matched the number of recording sessions. Within each batch, duplicate session-channel pairs were removed. For each batch, we computed the total similarity matrix as described above. The total similarities from all batches formed a null distribution representing the similarity between unrelated neurons. The threshold similarity score was then defined as the 95th percentile of this null distribution, ensuring a false positive rate of 5%. This procedure was repeated over 200 iterations to obtain a robust threshold estimate, resulting in subject-specific similarity cutoffs (Monkey A: 0.554; Monkey B: 0.563).

### Clustering tracked neurons

To determine neuron identities across sessions, we used total similarity matrices to group neurons into clusters. For each MW, similarity scores with all other units were compared to the predefined threshold. Two clustering approaches were considered. The first approach considered was strict clustering, which requires that all units in a group have similarity scores exceeding the threshold with every other unit in the group. The alternative approach considered was flexible clustering, which allows a unit to be included if it meets the similarity threshold with at least one other unit in the group. We adopted the flexible clustering approach, as it accounts for gradual waveform changes over time that can improve tracking accuracy. A depth-first search algorithm (43) was implemented to iteratively group units based on these criteria. To resolve potential conflicts—such as multiple units from the same channel and session being grouped together—the unit with the highest average similarity score was retained, while the others were excluded. This step minimized false positives and ensured biologically plausible tracking.

### Glossary for neuron tracking

#### Sorted neurons

a collection of neurons that were sorted online by the Grapevine NIP system in the beginning of a session. Spiking activities were recorded for all the sorted neurons, including spike times and local field potentials (LFPs). The Grapevine NIP system allows at most four sources of neural activities to be sorted from one channel, yielding at most four sorted neurons in a single channel. In our tracking algorithm, if multiple neurons were sorted in a channel from a session, those sorted neurons were treated anonymously with other sorted neurons in other sessions.

#### Mean waveforms (MWs)

Our tracking algorithm relied on the waveform shapes that were averaged over all the spiking instances in a sorted neuron. Mean waveforms were coined to clarify that it was not the individual spike waveform from each spiking instance that were used to track neurons.

#### Tracked neurons

A collection of sorted neurons resulting from the tracking algorithm. If a number of sorted neurons from multiple sessions were grouped together by the algorithm, they were regarded as the same sorted neuron that were tracked longitudinally, and hence a tracked neuron.

#### Cluster (v.)

The action of grouping MWs corresponding to sorted neurons from different sessions together. Clustering was referred to as the process of tracking neurons across sessions.

### Spiking properties

The mean firing rate of each sorted neuron was calculated as the total number of spikes divided by the session duration. To assess the stability of spiking properties, we examined the peak amplitude, trough amplitude, and mean firing rate across sessions. Stability was quantified by computing the relative change in each metric, defined as the difference between the first and last session, normalized by the value in the first session. To statistically evaluate stability, we performed two-tailed one-sample t-tests on the relative changes. The null hypothesis was that the relative change was not significantly different from zero, indicating no systematic variation over time. Since the number of sessions varied widely across neurons (ranging from three to approximately 30), this analysis was restricted to the first and last session of each tracked neuron.

Inter-spike intervals (ISI) were aggregated across all spike events in each session. To visualize ISI distributions, we applied kernel density estimation (KDE) using the *kdeplot* function from the seaborn library. The coefficient of variation (CV), defined as the ratio of the standard deviation to the mean of the ISI distribution, was used to quantify variability in spiking patterns (44). To assess ISI stability over time, we compared CVs across sessions within each tracked neuron. A Mann-Whitney U test was used to compare each session’s CV against the first session’s CV. If the mean CV across all sessions differed significantly from the first session (p < 0.05), we rejected the null hypothesis of stability and classified the neuron as having unstable ISI. If the difference was not statistically significant, the neuron was classified as stable.

### Neural tuning properties

After tracking neurons across sessions, we assessed the stability of their directional tuning properties. To ensure meaningful comparisons, only neurons appearing in at least three sessions were included in this analysis.

Tuning properties were quantified using a cosine tuning model, which describes the relationship between firing rate and movement direction (45). Firing rates were measured within a 300 msec window aligned to movement onset and were calculated as the number of spikes in that period divided by the duration. The following equation was used to fit tuning curves, where *y* represents the firing rate, *θ* is the movement direction, *b*_0_ is the baseline firing rate, *b*_1_(modulation depth, MD) quantifies tuning strength, and *θ*^∗^(preferred direction, PD) indicates the movement direction eliciting the highest firing rate.

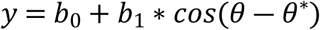

To assess PD stability, we calculated PD span, defined as the minimal angular range covering all observed PDs for a given neuron. To test whether observed PD spans were significantly small, we formulated a null hypothesis that PDs were randomly drawn from a uniform distribution spanning 0° to 360°. A null PD span distribution was generated by sampling k random PDs from a uniform distribution (where k was the number of recorded sessions for that neuron) and computing the span in each iteration. This process was repeated 100,000 times to construct a null distribution. If the observed PD span was smaller than the 5^th^ percentile of the null distribution, the neuron was classified as having a significantly stable PD at the p<0.05 significance level.

### Spike-field analyses

To assess the relationship between spiking activity and local network dynamics, we analyzed spike-triggered averaging (STA) of LFPs and spike-field coherence (SFC).

STA was used to examine the average LFP waveform aligned to spike times, providing insight into how neuronal firing correlates with local oscillatory activity. A well-defined STA waveform suggests that spikes tend to occur at specific LFP phases, indicating functional coupling between single-neuron activity and network oscillations (46,47). To evaluate STA stability, we quantified two key features: (1) phase of the STA waveform, meaning the preferred phase at which spikes occurred relative to the LFP, and (2) peak-to-trough amplitude, which was the magnitude of STA deflection, normalized to the first session for comparability across tracked neurons. To assess whether these features remained stable across sessions, we constructed a bivariate distribution of phase differences and normalized amplitudes for all tracked neurons. A Hotelling’s T-squared test was used to determine whether these distributions deviated significantly from the first session using a standard significance level of 0.05.

SFC was computed to quantify the degree to which neurons were phase-locked to oscillatory LFP components. This metric assesses how consistently spikes occur at specific phases of LFP oscillations across trials (48). The SFC was analyzed in different frequency bands, with a particular focus on theta-band coherence (2-6 Hz), as theta oscillations have been implicated in sensorimotor network (36). To assess the stability of SFC, we computed the cross correlation between the SFC in the first session of each tracked neuron and its SFC in subsequent sessions. This approach accounts for slight temporal shifts in SFC peaks over time. To determine whether the observed time lags were significantly different from zero, we performed a Wilcoxon signed-rank test for each tracked neuron. Additionally, we pooled the maximum correlation values from each cross-correlation analysis to evaluate population-level SFC stability.

By analyzing both STA and SFC across multiple recording sessions, we assessed the stability of spike-field interactions over time. This allowed us to determine whether neurons maintained their phase relationships with LFP activity despite potential changes in firing properties.

### Statistical Analysis

Statistical analyses were performed using custom written Python3 scripts and a significance level set at 0.05. Effect sizes were computed using Cohen’s D (Cohen, 1988). Two-tailed one-sample t-test, Hotelling’s T-squared test, and Mann-Whitney U tests were used for statistical comparisons. No statistical tests were run to predetermine the sample size.

## Results

### Neuron tracking

Two rhesus macaques (Monkeys A and B) implanted with chronic microelectrode arrays (MEAs) in the primary motor cortex (M1) and premotor cortex (PMd) were trained to perform a center-out brain-machine interface (BMI) task (see Methods). Spiking activity from M1/PMd was sorted online using the hoop method (39) in the Grapevine NIP system. A subset of online sorted units was selected prior to each session to control cursor velocity via a Kalman filter decoder (Fig 1A).

In each trial, the subject initiated movement from a center target toward one of eight peripheral targets under time constraints (center target hold: 200 msec, peripheral target hold: 200 msec, total movement time limit: 10 s) to obtain a juice reward. Each session consisted of 336 successful trials (42 per target direction). The number of recorded neurons increased over the initial sessions as the MEAs stabilized (Fig 1B). A maximum of 53 and 140 neurons were recorded in a single session for Monkeys A and B, respectively. By the final session, approximately 50% of the maximal number of neurons was captured, likely due to changes at the electrode-tissue interfaces, such as electrode drift (49) and glial scarring (50).

To investigate the longitudinal stability of neuronal activity, we tracked sorted neurons within the same channel across sessions based on their mean waveforms (MWs). Given that electrode contacts were sufficiently spaced apart to prevent neurons from migrating across channels, tracking was performed independently within each channel (Fig 1C). We computed pairwise total similarities between MWs from different sessions, defined as the mean of Fisher-transformed Pearson correlations and log-transformed Euclidean distances, each rescaled between 0 and 1. Tracked neurons were identified by grouping together subsets of MWs if at least one MW exceeded a similarity threshold (see Methods).

Given that waveform shapes may gradually change over time (51), we applied a flexible clustering criterion, where an MW needed to surpass the threshold with at least one other MW in the group. This approach allowed for minor waveform variations. Potential conflicts arose when multiple sorted neurons from the same channel were grouped together. To resolve these conflicts, only the sorted neuron corresponding to the MW with the highest average similarity score to the rest of the cluster was retained, ensuring biologically plausible tracking.

After applying the tracking algorithm, we identified 116 tracked neurons (from 656 MWs) in Monkey A and 354 tracked neurons (from 1,774 MWs) in Monkey B (Fig 2A). To ensure reliability in subsequent analyses, we excluded poorly online sorted neurons and any tracked neurons that appeared in two or fewer sessions, resulting in the removal of 60.4% (Monkey A) and 67.4% (Monkey B) of all MWs. This filtering prioritized robustly tracked neurons for assessing stability in firing properties and tuning. The number of sessions each neuron was tracked varied widely (Fig 2B). The distribution of tracking durations was right-skewed, meaning most neurons were tracked for relatively short periods, but some were tracked for over 100 days (Monkey A: 112 days, Monkey B: 103 days). The maximum number of sessions in a single tracked neuron was 23 and 32 for Monkeys A and B, respectively.

**Fig 2.**
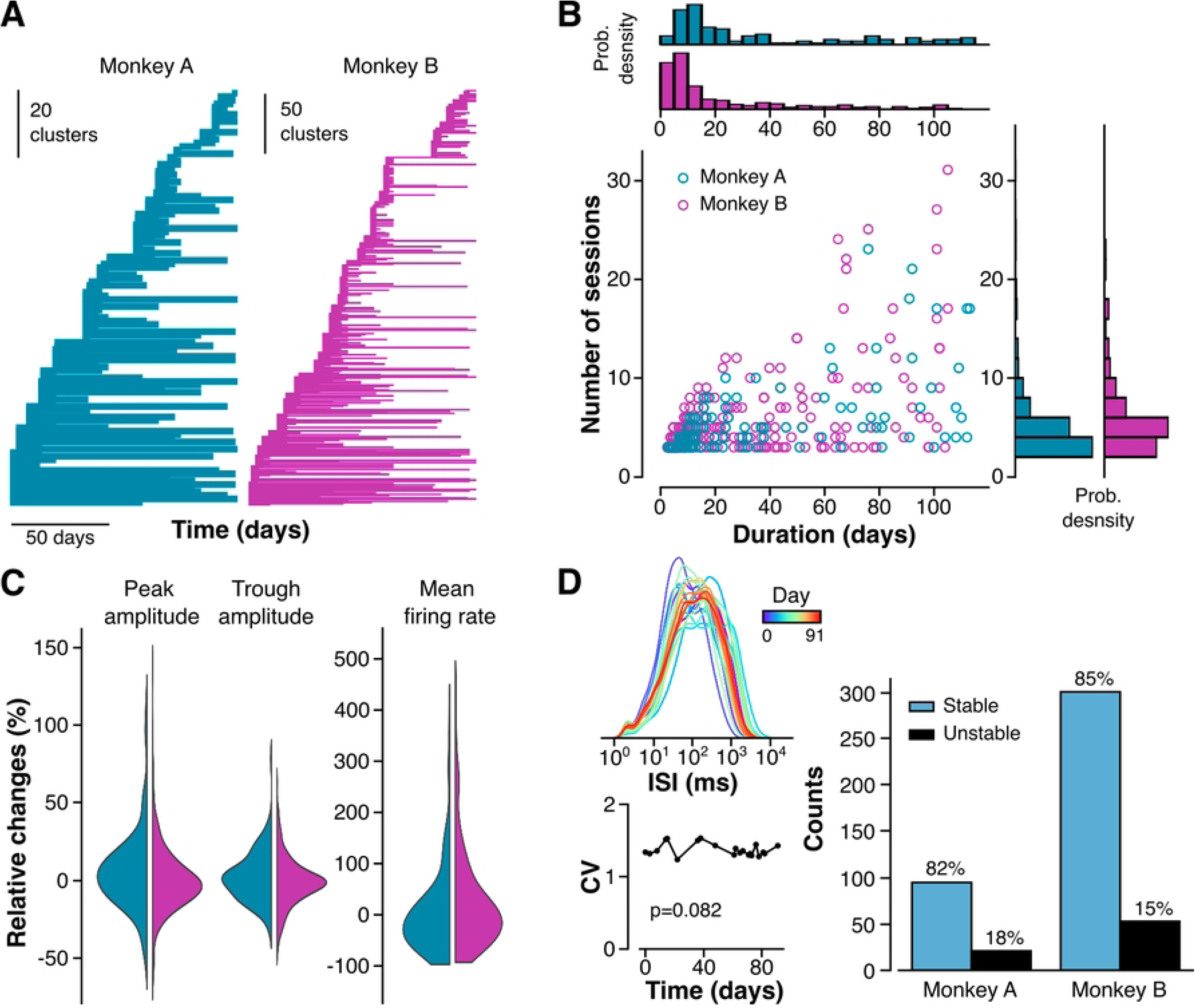
Tracking results and spiking properties. (A) The durations of tracked neurons sorted by their first appearance. (B) The number of sessions and duration of each tracked neuron, as represented as a circle in the Fig. Marginal distributions of each variable were plotted on the top and the right. (C) The relative changes in peak amplitudes, trough amplitudes, and mean firing rates (Monkey A: peak amplitudes, p<0.05, Cohen’s D=0.215; trough amplitudes, p>0.05. Monkey B: peak and trough amplitudes, p>0.05). (D) All ISI distributions from all the sessions in an example tracked neuron that spanned across 91 days (top left). The coefficient of variations for each session was plotted against the time of recording (bottom left). In this example, the CV samples were not significantly different from the first CV (N=21, p=0.082). The numbers and percentages of tracked neurons that had stable and unstable ISI (right).

### Spiking properties of tracked neurons

Since tracking was based on MWs, we first verified waveform stability across sessions (Fig 2C, left). We quantified peak and trough amplitude changes between the first and last session of each tracked neuron, normalizing differences by the first session’s amplitude. Most waveform features remained unchanged (Monkey A, trough amplitudes: p > 0.05; Monkey B, peak and trough amplitudes: p > 0.05; two-sided one-sample t-tests), with only a minor change in peak amplitude for Monkey A (p < 0.05, Cohen’s D = 0.215). These results confirm that our algorithm effectively tracked neurons with stable waveforms.

We next analyzed mean firing rates, calculated over entire sessions (Figure 2C, right). Although firing rates showed greater variation than waveform amplitudes, most changes were centered around zero, indicating only minor session-to-session fluctuations. Monkey B exhibited significant changes in mean firing rates, whereas Monkey A did not. This variability is expected, as firing rates were not used in the tracking algorithm.

We further examined inter-spike interval (ISI) distributions over time (Fig 2D, left). While some tracked neurons maintained stable ISI distributions across sessions, others exhibited shifts in spiking patterns. To quantify ISI stability, we calculated the coefficient of variation (CV) for each session and compared it to the first session in which the tracked neuron appeared using Mann-Whitney U tests. If the CV differed significantly (p < 0.05), the neuron was classified as unstable; otherwise, it was labeled stable. We found that 82% (Monkey A) and 85% (Monkey B) of tracked neurons had stable ISI distributions (Fig 2D, right), consistent with the firing rate analysis. Notably, we found multiple tracked neurons with drifting ISI distributions (Fig 3), which were recorded for more than 60 days.

**Fig 3.**
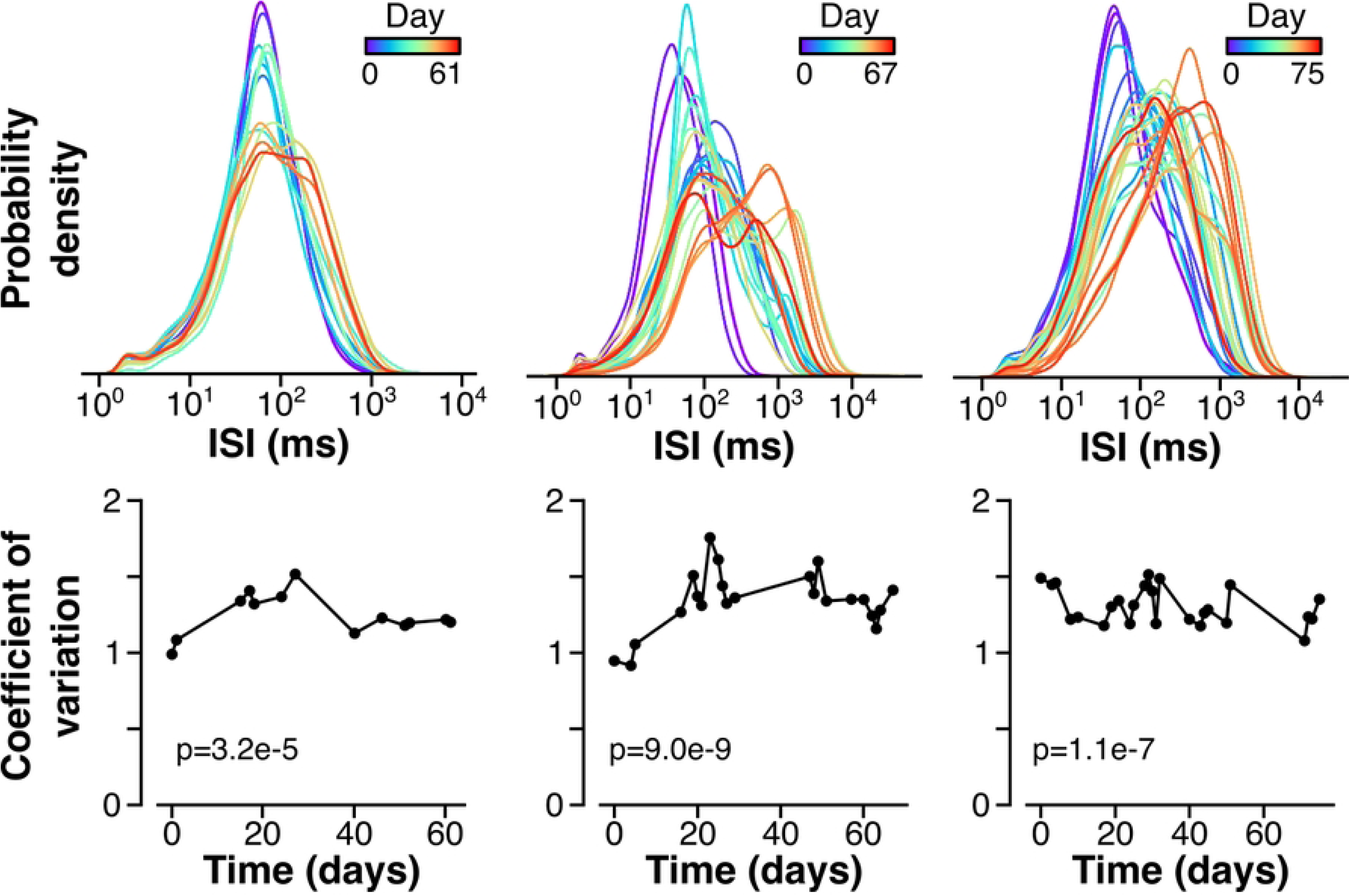
Drifting inter-spike intervals (ISI). Examples of unstable ISI color-coded with their times of recordings (top row) and their corresponding coefficients of variations (CV) across time (bottom row). Mann-Whitney U tests were used to test the CVs against their first samples (left: N=13, p=3.2e-5; center: N=22, p=9.0e-9; right: N=25, p=1.1e-7).

### Preferred directions of tracked neurons

To assess whether tuning properties remained stable across sessions, we fit a cosine tuning model (45) to each tracked neuron using the firing rate *y* and the target direction *θ* by the following equation, where the fitted parameters *b*_0_, *b*_1_, and *θ*^∗^ denote the baseline firing rate, modulation depth, and the preferred direction (PD), respectively.

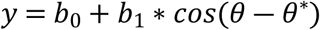

We first examined PD changes between neighboring sessions (Fig 4A). At the population level, PD shifts were not significant, suggesting that PDs remained stable over time. However, this analysis only compared session-to-session variations rather than long-term consistency.

**Fig 4.**
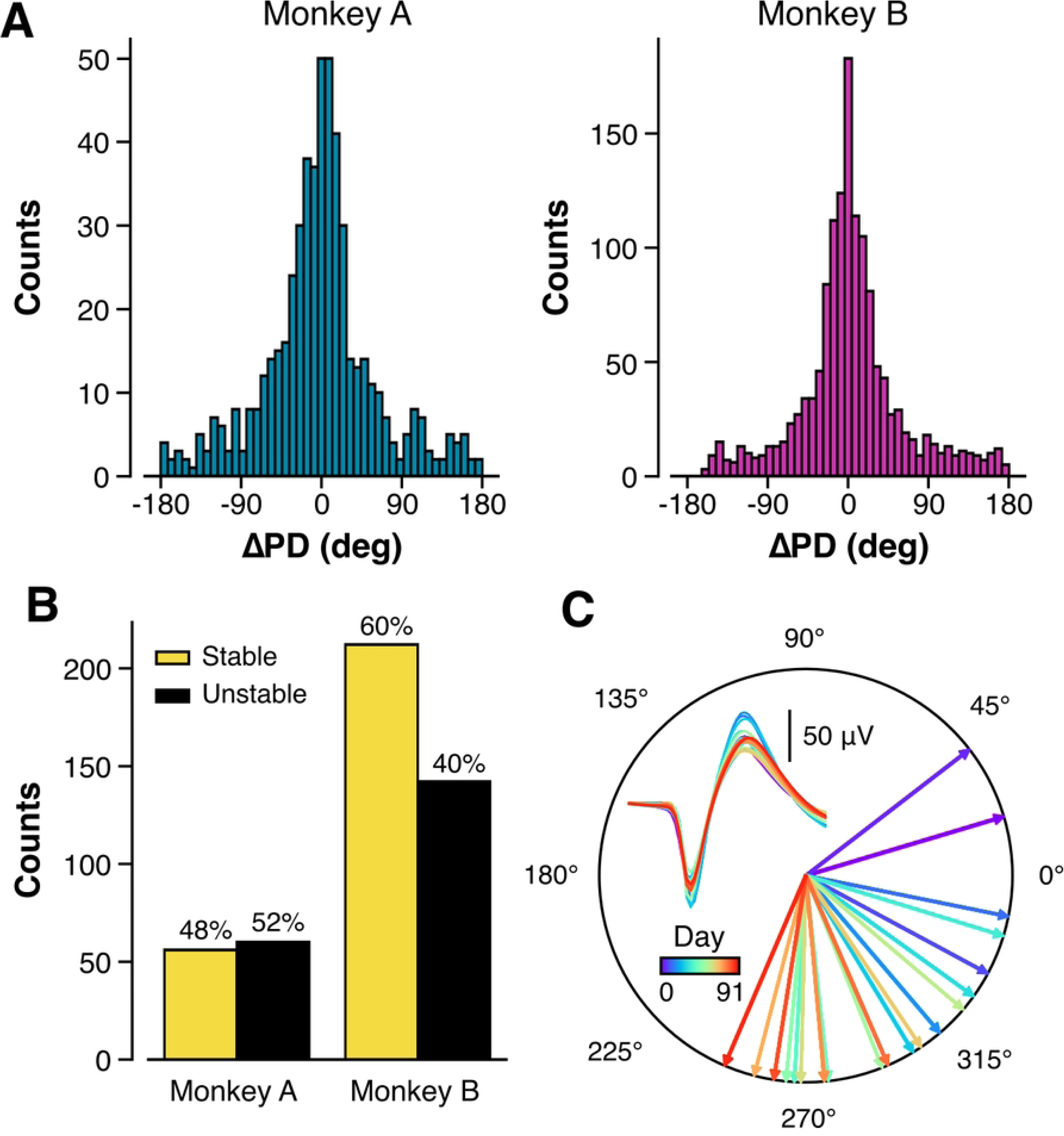
Preferred direction (PD) analysis. (A) The changes in PDs between neighboring sessions in tracked neurons (Monkey A: N=540, p=0.54; Monkey B: N=1420, p=0.36. One-sample t-test). (B) The numbers and percentages of tracked neurons that had stable and unstable PDs, as characterized by the PD span. (C) The PDs and their corresponding mean waveforms of an example tracked neuron.

To assess PD stability across all sessions in a tracked neuron, we computed the minimal angular range encompassing all PDs and refer to this as PD span. If a tracked neuron’s PDs remained consistent, its PD span would be small. To evaluate statistical significance, we formulated a null hypothesis assuming PDs were randomly distributed from 0° to 360°. We then generated 2,000 surrogate PD spans from this uniform distribution to construct a null distribution. The 5th percentile of this distribution served as the threshold for significant PD stability.

We found that 48% (Monkey A) and 60% (Monkey B) of tracked neurons had stable directional tuning (Fig 4B). An example tracked neuron (Fig 4C) exhibited a PD around 37.6°, gradually shifting clockwise to 246.8° over 91 days, yet its PD span (150.8°) remained well below the null threshold (266.4° for 21 sessions), demonstrating remarkable stability.

### Spike-field relationship

Having established that tracked neurons can exhibit stability in spiking properties (waveforms, firing rates, and ISI) and directional tuning, we next examined whether spike-field relationships remained stable across sessions. Local field potentials (LFPs) are considered more stable over time relative to single-unit activity, as individual neurons may become undetectable due to electrode drift or other factors. However, it remains unclear whether spike-LFP interactions remain stable when neurons are reliably recorded across multiple sessions.

To investigate this, we first computed spike-triggered averages (STA) of LFP signals for each tracked neuron across sessions. We observed clear oscillatory patterns in the STA, indicating that spikes were phase-locked to specific LFP fluctuations. This trend persisted across multiple sessions in most tracked neurons, suggesting that their coupling with local oscillatory activity remained stable over time (Fig 5A). To quantify STA stability, we pooled all tracked sessions for each subject and computed phase differences and normalized log-scaled amplitudes relative to the first session of each tracked neuron. No significant changes in STA amplitude or phase distributions over time were observed in either subject (Fig 5B. Monkey A: F(2,610) = 0.799, p = 0.45; Monkey B: F(2,1710) = 2.364, p = 0.09; Hotelling’s T-squared test), indicating that spike-phase relationships were maintained across sessions.

**Fig 5.**
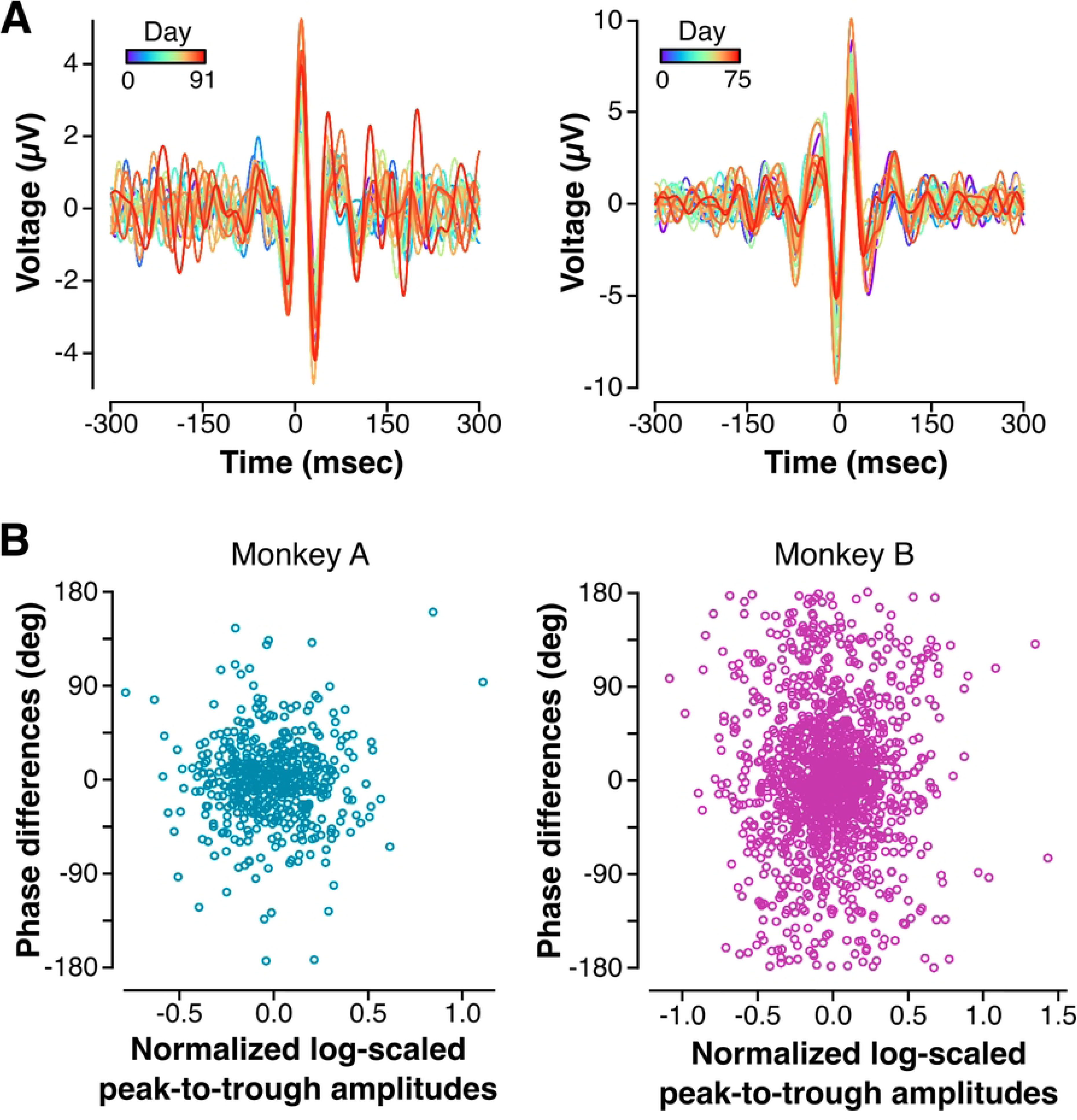
The spike-triggered averages (STA) of LFPs. (A) STA of example tracked neurons. (B) The distributions of phase differences and normalized log-scaled peak-to-trough amplitudes across all sessions in all tracked neurons in Monkey A (left, F(2,610)=0.799, p=0.45; Hotelling’s T-squared test) and Monkey B (right, F(2,1710)=2.364, p=0.09).

In addition to STA, we assessed spike-field coherence (SFC) to measure the consistency of spike timing relative to oscillatory LFP activity around certain behavioral metrics. SFC was computed around movement onset, using LFP signals recorded from the same channel as the spiking unit. To facilitate visualization across multiple sessions, we focused on theta-band coherence (2–6 Hz), which has been reported to play a significant role in sensorimotor network (36). We observed a strong increase in theta-band coherence around movement onset (Fig 6A), suggesting that spiking activity was temporally coordinated with theta oscillations during task execution. The SFCs remained highly consistent across the nine-day recording period.

**Fig 6.**
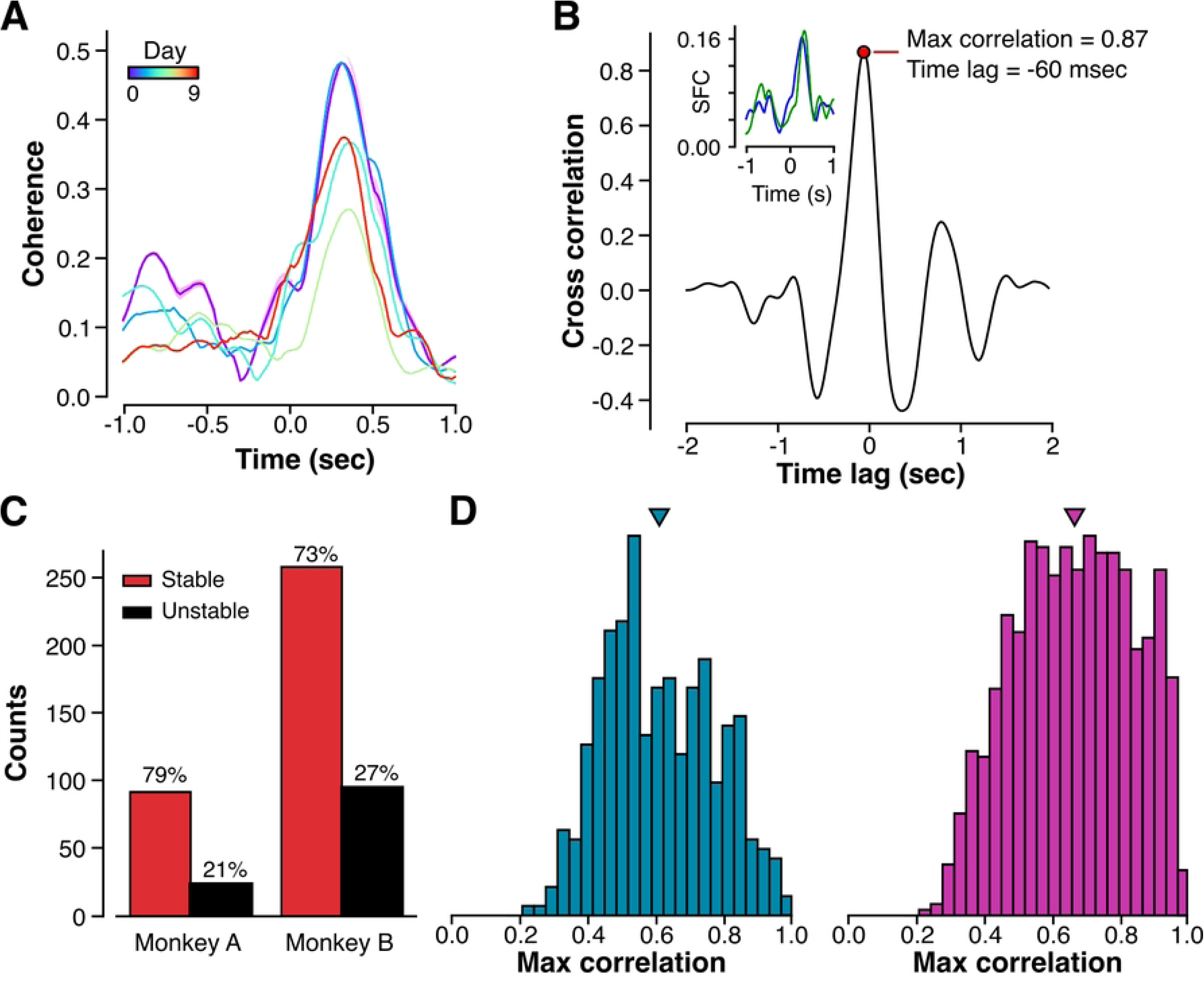
Spike-field coherence (SFC) analysis. (A) SFC of an example tracked neuron. (B) The cross-correlation analysis from an example pair of SFC time series (inset). (C) The numbers and percentages of tracked neurons that had stable and unstable SFC, as characterized by the statistical difference of time lags in each tracked neuron. (D) The probability density of maximum correlation values in all sessions across all tracked neurons for Monkey A (left) and Monkey B (right). Triangles pointed to the mean of the population level of maximum correlation (Monkey A: 0.61; Monkey B: 0.66).

To quantify the stability of SFC across sessions in a tracked neuron, we computed cross-correlation between the SFC from the first session and subsequent sessions, accounting for potential temporal shifts in SFC peaks over time. In an example pair of SFCs (Fig 6B), cross-correlation analysis yielded a maximum correlation of 0.87 with a time lag of - 60 msec, indicating that the SFC time series were highly similar when aligned with this shift. For each tracked neuron, we applied the Wilcoxon signed-rank test to determine whether the distribution of time lags across sessions significantly deviated from zero. If the test indicated a significant difference, the neuron was classified as having unstable SFC.

Overall, 79% (Monkey A) and 73% (Monkey B) of tracked neurons exhibited stable SFCs (Fig 6C). Moreover, we collected the maximum correlation values from each cross-correlation analysis and found that the population-level maximum correlation was 0.61 in Monkey A and 0.66 in Monkey B (Fig 6D, triangles). For both subjects, over 90% of the maximum correlations exceeded 0.4 (Monkey A: 91.58%; Monkey B: 92.57%). These results indicate that SFC remained highly correlated with its first-session values, and the majority of tracked neurons did not exhibit significant temporal shifts in SFC across sessions.

Finally, we investigated the proportions of tracked neurons that exhibited stable functional properties, such as ISI and PD, as well as stable relationships with local dynamics, such as SFC. Across both subjects, a great portion of tracked neurons are stable in all functional properties and spike-field relationships (32% for Monkey A and 36% for Monkey B), while almost all tracked neurons are stable in at least one property (115 out of 116 tracked neurons in Monkey A and 348 out of 354 tracked neurons in Monkey B). Other combinations also show similar percentages of tracked neurons across subjects that maintained highly consistent functional properties over time and that had variable “plastic” properties over time (Fig 7).

**Fig 7.**
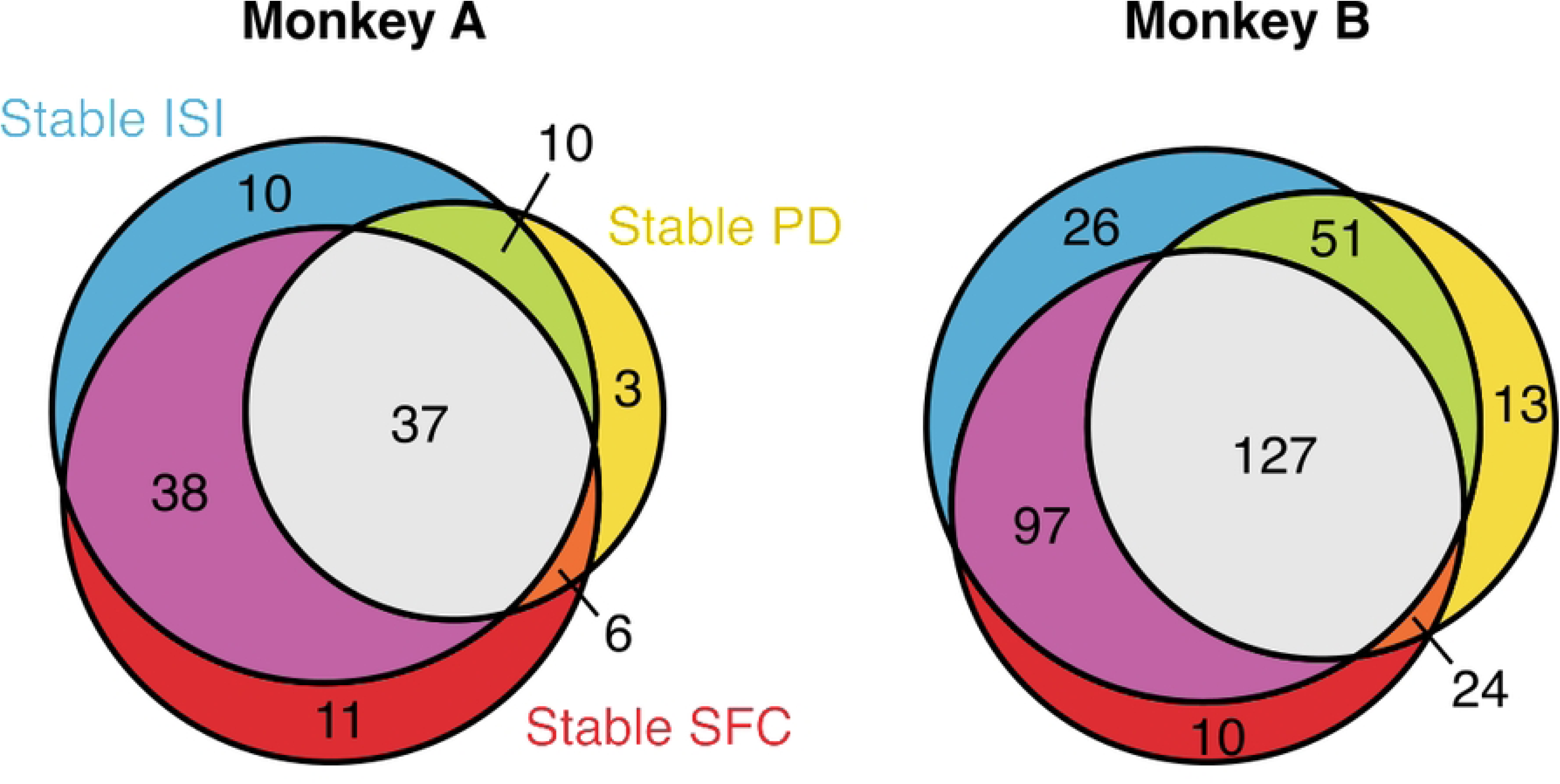
Venn diagram of the number of stable neurons. The Venn diagrams showed the numbers of tracked neurons in each factorial combination of the stability of ISI, PD, and SFC. Almost all of tracked neurons were stable in at least one functional property (115 out of 116 tracked neurons in Monkey A and 348 out of 354 tracked neurons in Monkey B), whereas more than 30% of tracked neurons have stable ISI, PD, and SFC (32% for Monkey A and 36% for Monkey B).

## Discussion

The ability to track individual neurons over extended periods is critical for understanding neural stability and plasticity. In this study, we employed a waveform-based tracking approach to identify and track neurons across more than 30 sessions acquired during a three- to four-month time period. We demonstrated that spiking dynamics, directional tuning, and spike-field interactions remained stable over time a large proportion of neurons, while the remaining neurons were more plastic. Longitudinal neuron tracking provides insights into the persistence of neural representations and has direct applications in developing more robust brain-machine interfaces (BMIs) and studying learning-related neurophysiological changes.

Unlike tracking methods optimized for high-density electrode arrays or tetrodes, our approach is specifically designed for conventional MEAs, where inter-electrode distances prevent neurons from being detected across multiple channels. While our method leverages mean waveforms to identify the same neurons across sessions, other approaches, such as UnitMatch (29), incorporate spatial information, making them more suited for high-density recordings. The trade-off is that our method prioritizes a more computationally efficient implementation for standard MEAs while maintaining high specificity by controlling false positives. However, as with all tracking methods, challenges remain in distinguishing between gradual waveform drift and true neuronal turnover.

Neuron tracking is inherently a balance between sensitivity and specificity. In this study, we prioritized minimizing false positives by setting a similarity threshold, ensuring that neurons identified as “matches” were highly likely to be the same. However, this approach does not explicitly control false negatives, meaning that some neurons that persisted across sessions may have been excluded due to waveform variability (27). Future tracking methods could integrate supervised learning techniques trained on known same and different neurons to better optimize this trade-off potentially from simultaneous two-photon imaging with electrophysiology (52,53).

A limitation of our algorithm is its computational scalability. Since each neuron must be compared to all previously recorded neurons in a given channel, our implementation scales quadratically (O(n²)). While feasible for arrays with standard channel counts (typically < 256 channels), this could pose challenges for high-density arrays where thousands of units are recorded simultaneously. Optimizing the tracking process using clustering-based pre-filtering or dimensionality reduction techniques may improve computational scalability without sacrificing accuracy. Another important consideration is the duration of the experiment. Our recordings spanned 34–44 sessions across 105-113 days and may not generalize to longer time periods of several months to years. Additionally, we excluded over 60% of sorted neurons due to strict tracking criteria, which could have led to an underestimation of neural variability. However, these limitations highlight again the importance of a reliable tracking algorithm that can identify neurons accurately over the long-time horizons.

Our results demonstrated that tracked neurons exhibited stable waveform properties over time, which is expected given that waveform similarity was the primary criterion for tracking. Traditionally, tracking method rely on the assumption that waveform characteristics remain stable over time, which may not always hold (28). Our method addresses this limitation by allowing waveform similarities to evolve across sessions, accommodating gradual changes while maintaining reliable tracking. Besides waveform stability, we also observed that ISI distributions remained largely unchanged for most neurons, consistent with prior studies suggesting stable neuronal firing patterns over extended periods (3). This stability suggests that fundamental aspects of neuronal excitability and synaptic inputs remain relatively constant despite ongoing network plasticity. While ISI remained stable in most neurons, some tracking methods assumed that ISI distributions are intrinsically stable and use them as a primary feature for neuron tracking (26). This could potentially reflect the limited two-week duration of their recordings (26), where shifting in ISI distributions were negligible. Our results challenge this assumption by showing that ISI variability can occur over longer timescales. This highlights the importance of selecting appropriate tracking features based on the underlying stability of neural properties rather than assuming their stationarity across time. Directional tuning properties demonstrated both stability and plasticity across sessions. While session-to-session PD shifts were relatively small across tracked neurons, which indicated stable tuning at the population level, some neurons exhibited greater fluctuations in PD. This suggests that motor cortical neurons maintain directional selectivity over time while allowing for flexibility, which may reflect adaptive changes in response to learning or environmental shifts (54,55).

One important question is whether stability in tuning properties correlates with motor learning. If neurons with highly stable PDs contribute more to precise movement execution, this could suggest that stable tuning properties are advantageous for motor control. Conversely, neurons with greater PD variability may participate in adaptive learning, allowing the system to adjust to task demands over time. Further studies examining how PD stability relates to task performance and BMI control accuracy could provide deeper insights into the functional role of neural stability.

Beyond PD stability, spike-field interactions also remained stable in a subset of neurons, while others exhibited fluctuations in coherence over time. STA analysis revealed that spikes were not only phase-locked to the LFP signal but that this phase-locking persisted across multiple sessions. Similarly, SFC analysis further demonstrated that some neurons maintained consistent phase-locking with LFP oscillations that were also time-locked to behavior. Our findings were consistent with previous reports showing that movement toward a target elicited strong theta-band coherence (35,36). Furthermore, our analysis extends this understanding by demonstrating that this coherence can persist across multiple recording sessions.

Our findings may help reconcile discrepancies in previous studies on long-term neural stability, where some have reported stable neural dynamics (3,11–13), while others have observed significant fluctuations over time (14–17). Based on our analysis, these differences could arise from variations in neuron sampling across sessions or differences in the duration of recordings. Studies that report stability over short timescales may not capture the gradual changes that emerge over longer periods. Conversely, findings of instability may result from tracking inconsistencies rather than true neuronal plasticity. By implementing a tracking method that accounts for waveform evolution over time, our study provides a more comprehensive perspective on neural stability, highlighting both its persistence and its capacity for adaptation.

The ability to track neurons across sessions has several important applications in basic and clinical neuroscience. One immediate application is pooling neural data across multiple sessions to construct more stable neural representations. Previous studies (26,27) suggested that aggregating multi-session neural responses improves statistical power for decoding models, making tracking methods crucial for large-scale neural population analyses. Longitudinal tracking also provides a valuable framework for studying learning and adaptation. By monitoring the same neurons over time, researchers can assess how neural representations evolve during skill acquisition, motor adaptation, or rehabilitation. This is particularly relevant for studying motor plasticity and understanding how neural circuits reorganize following injury or disease. For BMI applications, tracking neurons can enhance decoder stability and calibration. A persistent challenge in BMI design is neural signal drift or loss, which necessitates frequent recalibration of decoders (56) that takes time every session (57) and may impede recruitment of neural plasticity (3). By identifying stable neurons over time, our approach could facilitate the development of adaptive BMIs that compensate for gradual neural changes, improving long-term performance and reducing patient effort in maintaining control. Finally, our findings could guide investigations into intrinsic neural variability, such as changes in ISI distributions that may reflect ion channel regulation or synaptic plasticity. Understanding the biological mechanisms underlying neural stability and plasticity could lead to new insights into the fundamental principles governing motor control and learning. Future work should aim to refine tracking methodologies, extend recording durations, and explore the relationship between neural stability and motor learning to further advance our understanding of cortical dynamics over time.

## Acknowledgments

This work was supported by Whitehall Foundation [2022-12-071] and National Science Foundation CAREER Award [2145412].

## Author Contributions

H.L.: Conceptualization, Methodology, Formal Analysis, Writing – Original Draft Preparation, Writing – Review & Editing

H.M.S.: Methodology, Writing – Review & Editing

Y.Z.: Methodology, Writing – Review & Editing

C.R.B.: Methodology, Writing – Review & Editing

E.C.H.: Methodology, Writing – Review & Editing

S.R.S.: Conceptualization, Methodology, Investigation, Writing – Review & Editing

